# Investigating Ecological Interactions in the Tumor Microenvironment using Joint Species Distribution Models for Point Patterns

**DOI:** 10.1101/2023.11.14.567108

**Authors:** Joel Eliason, Arvind Rao

**Affiliations:** Department of Computational Medicine and Bioinformatics, University of Michigan, USA

**Keywords:** tumor microenvironment, spatial point patterns, joint species distribution models

## Abstract

The tumor microenvironment (TME) is a complex and dynamic ecosystem that involves interactions between different cell types, such as cancer cells, immune cells, and stromal cells. These interactions can promote or inhibit tumor growth and affect response to therapy. Multitype Gibbs point process (MGPP) models are statistical models used to study the spatial distribution and interaction of different types of objects, such as the distribution of cell types in a tissue sample. Such models are potentially useful for investigating the spatial relationships between different cell types in the tumor microenvironment, but so far studies of the TME using cell-resolution imaging have been largely limited to spatial descriptive statistics. However, MGPP models have many advantages over descriptive statistics, such as uncertainty quantification, incorporation of multiple covariates and the ability to make predictions. In this paper, we describe and apply a previously developed MGPP method, the *saturated pairwise interaction Gibbs point process model*, to a publicly available multiplexed imaging dataset obtained from colorectal cancer patients. Importantly, we show how these methods can be used as joint species distribution models (JSDMs) to precisely frame and answer many relevant questions related to the ecology of the tumor microenvironment.

## 1. INTRODUCTION

Similar to natural ecosystems, the TME comprises a dynamic web of relationships, where cancer cells, immune populations, stromal cells, and various molecular cues engage in complex communication and interdependence [1, 2]. Furthermore, just as environmental conditions influence species behavior in ecological systems, the TME’s microenvironmental factors, including nutrient availability, oxygen levels, and inflammatory signals, play a pivotal role in shaping cellular behaviors, proliferation, migration, and responses to therapy [3–5]. These interactions exert pivotal influences on tumor progression and therapeutic outcomes [6–9]. This ecosystem-like framework underscores the TME’s complexity and underscores the need to employ ecological modeling approaches to gain deeper insights into its intricate dynamics and therapeutic implications [10–14].

The advent of highly multiplexed imaging technologies that allow for the simultaneous measurement of dozens of cell-surface protein biomarkers in tissue samples has allowed for the creation of datasets that have precise spatial coordinates and cell-type annotations for each cell within a tissue [15]. Such multiplexed imaging technologies are of particular relevance to studies of the tumor microenvironment, in which multiplexed images can be used to ascertain the complex interrelationships between cell types and cell states [16, 17]. Such imaging technologies are similar to the remote sensing technologies employed in ecological studies, which allow the localization of various species and environmental covariates in space.

Spatial descriptive statistics are fundamental tools employed in such ecological studies to gain insights into the spatial patterns and relationships within natural systems. These statistics provide a quantitative framework for characterizing the arrangement, dispersion, and clustering of species or individuals across a landscape. Metrics such as nearest neighbor distances, spatial autocorrelation, and Ripley’s K-function are commonly used to assess the degree of spatial aggregation or dispersion of species within a given habitat [18–21]. By analyzing these patterns, ecologists can infer crucial information about species interactions, habitat preferences, and the influence of environmental factors on distribution. Such statistics have also been widely adopted by researchers investigating the tumor microenvironment [22–24]. More bespoke methods have also been developed, such as those in which a “neighborhood composition” vector is calculated for each cell [16, 25–27]. Other relatively simple statistics, such as quantifying the amount of immune infiltration based on the Immunoscore metric, in various regions of the TME, are also widely used in clinical models [28, 29].

While spatial descriptive statistics offer essential preliminary insights into ecological spatial patterns, more advanced techniques, such as Joint Species Distribution Models (JS-DMs), extend this analysis by capturing the complex interactions and dependencies that characterize ecological systems in a more holistic manner [30–34]. In particular, JS-DMs are able to model the spatial distributions of multiple species simultaneously, while accounting for the associations between each of these species, as well as their association with environmental variables. Within the context of the TME, these models can enable researchers to identify cell type-cell type or cell type-environment associations while adjusting for other associations, predict cell type distributions, and quantify the uncertainty of model parameters. Such capabilities are typically not available for spatial descriptive statistics.

Point process models are another widely used tool for modeling ecological relationships [21, 35–39]. Such models make explicit the probabilistic data-generating process that is assumed to generate an observed pattern of points, and thus they are amenable to uncertainty quantification and producing predictions [18]. A particular class of point process model, called multitype Gibbs point process (MGPP) models, are able to model the pairwise cell type interactions jointly [18, 40–42], along with cell type/environment interactions and thus can be used as JSDMs in the context of the TME ecosystem. Gibbs point process models explicitly model the attraction or inhibition between different points in space; MGPPs generalize this to when there are several types of points in space, such as different cell types [18].

More specifically, modeling interactions via MGPP models can improve on the aforementioned spatial descriptive statistics in a number of ways [18]: a) spatial variables, such as the concentration levels of various inflammatory factors, and non-spatial variables, such as organ of origin or patient demographics, can be easily adjusted for, allowing for the modeling of inhomogeneous distributions of cells; b) the joint modeling of pairwise interactions between cell types can be achieved, as opposed to the case of descriptive statistics, in which case only the marginal distribution of interactions can be estimated; c) uncertainty quantification of interaction effects can be performed; and d) predictions and residuals can be obtained from the model, allowing for the assessment of model fit, as well as the prediction of distributions of cells under new conditions.

## 2. METHODS

### 2.1. Dataset

The dataset used to showcase the MGPP method comprises multiplexed images of tumor tissue samples obtained from a publicly available repository for colorectal cancer [16]. The images capture different cell types within the tumor microenvironment, including cancer cells, immune cells, and stromal cells. In the original study, preprocessing was conducted on the images. This involved cell type classification and determination of spatial coordinates for each cell type.

Specifically, this dataset consists of 4 multiplexed images of 56 different cell-surface markers of tumor tissue sections from each of 35 CRC patients, with patients having varying degrees of disease progression and severity. These images were obtained via the PhenoCycler imaging technology (formerly known as CODEX) [17, 43]. The patients are divided into two groups, based on the histopathology of their tumors: those in which tertiary lymphoid structures (TLSs) are visible at the tumor invasive front, and those where diffuse inflammatory infiltration (DII) was present and no TLSs were visible. These patient groups will be subsequently referred to as CLR (Crohn’s-like reaction) and DII, with CLR patients generally having better survival prognosis [16]. 16 cell types have been annotated using the 56 different cell-surface markers and in total, the dataset consists of 244,504 annotated cells. In addition, patient metadata is included, including overall/progression-free survival times, tumor grade, and demographic variables such as age and sex. An example image can be seen in Figure 1.

**Figure 1:**
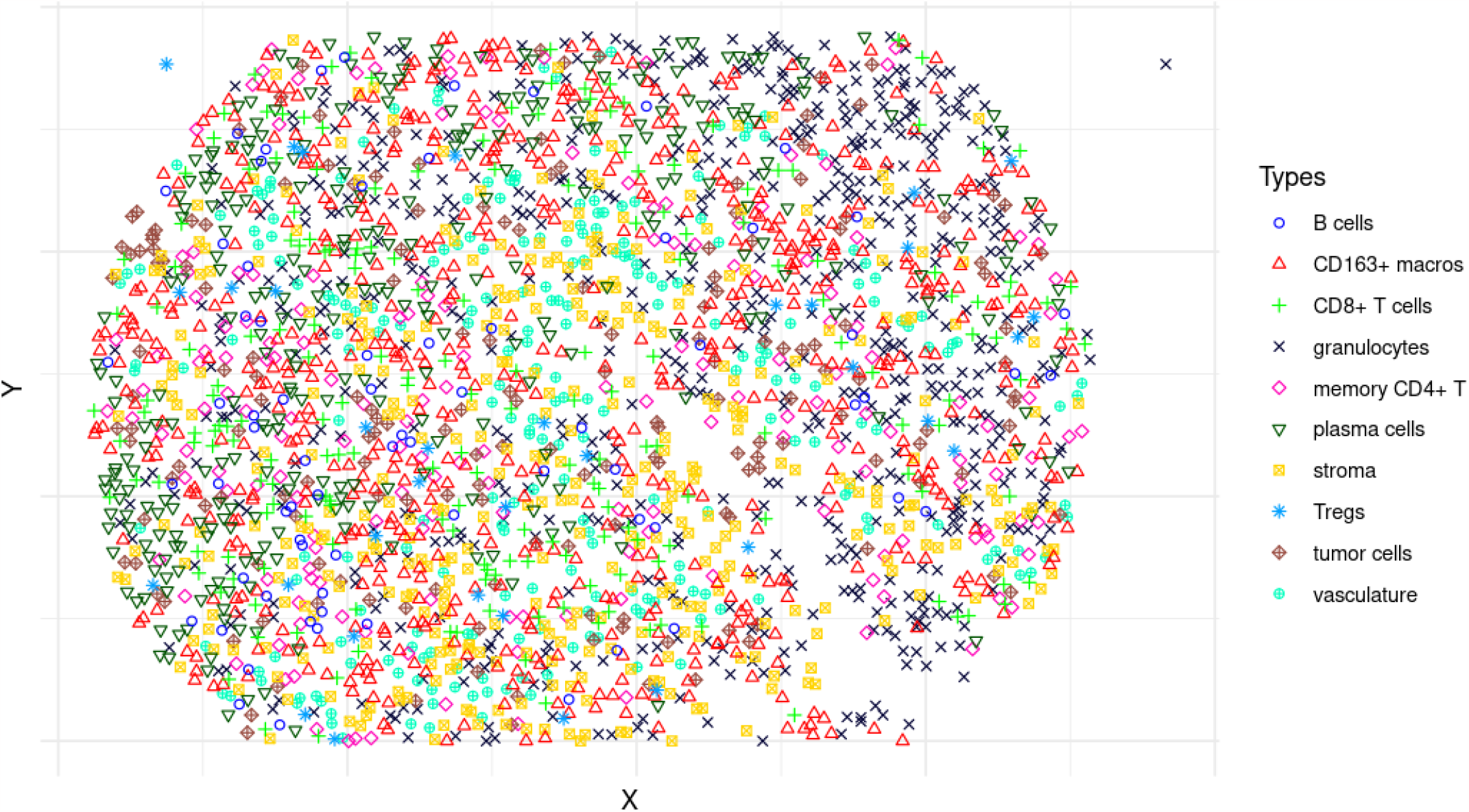
A point pattern representation of a multiplex image of a tissue section of the TME in a CRC patient.

### 2.2. Multitype Gibbs Point Process (MGPP) Models

Once cell types have been annotated, a multiplex image, such as in Figure 1, can be viewed as a multitype point pattern, or a realization **x** of a particular random variable **X** called a *multitype point process*. Subsequently, a MGPP model can be fit to this multitype point pattern. In the following, we will describe first a general *pairwise interaction* MGPP model, and subsequently the *saturated pairwise interaction* MGPP model.

Mathematically, a pairwise interaction MGPP model characterizes the probability density of a multitype point process **X** in the following way [40], with *i* and *j* indexing over the points in **x**:

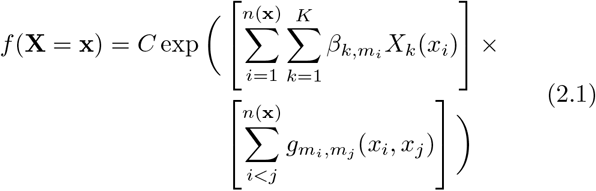

Here, *C* indicates a normalizing constant, while the first term in brackets signifies the “first-order” effects on the density of each cell type, that is, the effects from spatial and non-spatial variables. There are *K* such variables (the *X*_*k*_(*x*_*i*_)) per cell type. These variables are typically used to model “environmental variables”; in the context of the TME, this could include such variables as the concentration levels of various metabolic or inflammatory factors, or the distance to the nearest blood vessel. The second bracketed term contains the pairwise interaction functions 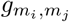(*x*_*i*_, *x*_*j*_), which are functions of two points *x*_*i*_ and *x*_*j*_ of types *m*_*i*_ and *m*_*j*_. These interaction functions are often parameterized with interaction parameters 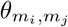, which typically quantify the direction and magnitude of interaction between points of type *m*_*i*_ and *m*_*j*_. For example, in the multitype Strauss model, the interaction function 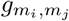 (*x*_*i*_, *x*_*j*_) is written as:

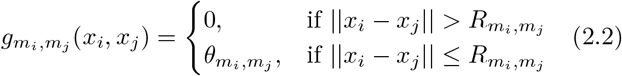

for user-defined *interaction radii* 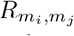 . The multitype Strauss model assumes that interactions between cells of types *m*_*i*_ and *m*_*j*_ do not exist if cells are farther than 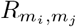 apart, and constant if they are within that radius.

One drawback of many pairwise interaction MGPP models, such as the multitype Strauss model, is that they are only able to model repulsion (and not clustering) between points - that is, that 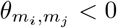 *<* 0. This has led to the development of MGPP models that do allow clustering, such as the *Geyer saturation* model [44] or the *saturated pairwise interaction* MGPP model [42]. For example, in the saturated pairwise interaction Gibbs point process, as in [42] and borrowing some of their notation, we have

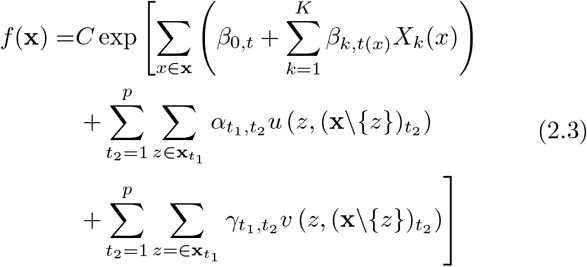

with

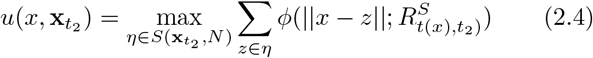

and *v* defined similarly as

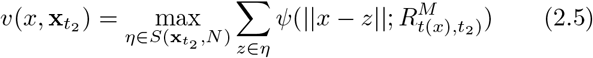

Here, *t*_*j*_ denotes the *j*^*th*^ cell type of *p* cell types, and 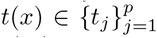denotes the type of point *x* ∈ **x**. Note that *t*(*x*_*i*_) = *m*_*i*_, as in our notation in Equation 2.1. Functions *φ* and represent user-defined short- and medium-range *potential functions* which set the shape of short- and medium-range interactions, and are dependent only on the distance between two points. *φ* and *Ψ* are commonly functions such as the exponential or step functions (Figure 2). 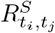 and 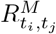 are user-defined short- and medium-range interaction radii, and the interaction parameters of interest, 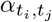 and 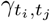, are interpreted as the strength and direction of interaction between any two types of points at short and medium ranges. 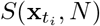 is the set of all subsets of points from 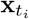 that are of size *N* - thus, *u* and *v* can be interpreted as the sum of the *N* largest potentials that point *x* has with points *z* ∈ **x** of type *t*_2_. Summing only the top *N* potentials (rather than all of them) is what leads to this being called a *saturated* MGPP model, and the parameter *N* is called the *saturation parameter* accordingly. Because of its great flexibility in being able to model clustering and repulsion, the saturated pairwise interaction MGPP model will be the one we will be using for the remainder of the paper as the JSDM.

**Figure 2:**
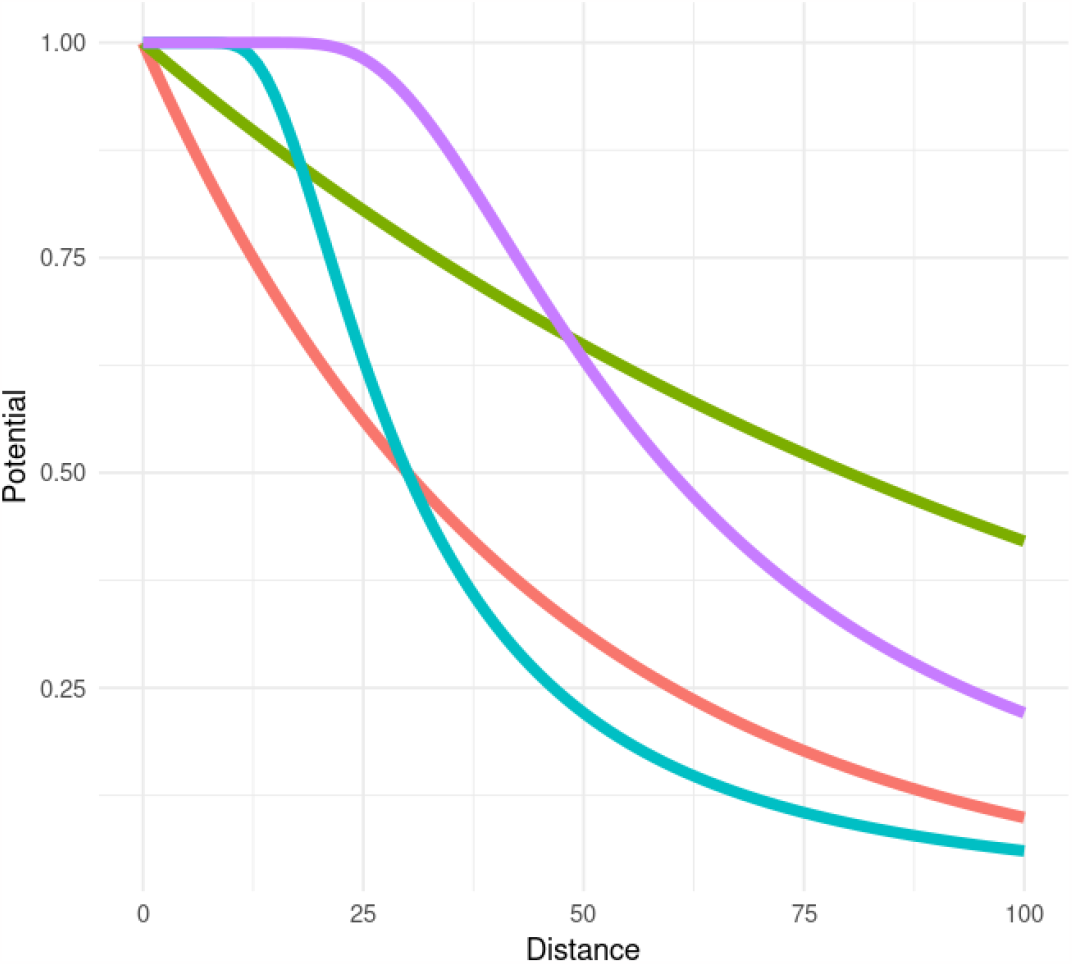
Examples of potential curves commonly used for *ϕ* and *ψ* in Equations 2.4 and 2.5.

Once the probability density *f* (**x**) of a MGPP model has been defined, estimating the parameters of this model can proceed via standard statistical techniques such as maximum likelihood estimation. However, because this is often quite computationally costly (due to the often intractable normalizing constant *C*), estimation techniques based on the *conditional intensity λ*(*y* | **x**) are often preferred. The *intensity λ*(*y*) of a point pattern at a location *y* is the expected number of points falling in a small window around *y*, divided by the area of that small window. The *conditional intensity* defined at a spatial location *y* is defined as 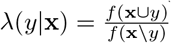, and can be interpreted as the “intensity at *y* given the rest of the spatial pattern **x**” [18]. Using the conditional intensity, we can then produce predictions from our MGPP model and subsequently assess model fit. For the specific formulation of the conditional intensity for the saturated pairwise interaction Gibbs point process model, see [42].

### 2.3. An Example Analysis

Here we demonstrate an example analysis to answer some of the questions related to the ecology of the TME. In this example, we jointly model the interactions between sufficiently abundant cell types in each image. Furthermore, we include as a spatial covariate the distance from each point to the nearest blood vessel, as a proxy for angiogenesis and oxygenation found in the TME. Specifically, we set out to answer the following:

1. Which cell types interact with one another, after adjusting for the interactions of other cell types and distance to blood vessels? Which cell types stay away from one another, again, after adjusting for the interactions of other cell types and the distance to blood vessels? What confidence can I have in individual parameter estimates?
  A. Do these interactions occur at close or far spatial scales?
  B. Which cell type most strongly uniquely interacts with tumor cells, on average, across the whole image?
2. Which cell types best predict the distribution of tumor cells at each location in space?
3. How much is the distribution of each cell type uniquely predicted by its distance to the nearest blood vessel?
4. What is the difference in interaction strength between each pair of cell types, comparing between the two patient groups (CLR and DII patients)?
5. What levels of intra- and inter-patient heterogeneity are there in how different cell types interact?
6. Are any of these parameter estimates useful clinically, for predicting patient outcomes?
7. How much can I trust the results from this model? i.e., how well does it predict spatial distributions of cells?

#### 2.3.1. Fitting MGPP models

We employed a systematic approach in fitting the MGPP models to the images from the CRC dataset. We focused our analysis on cell types that exhibited a minimum of 20 cells within a given image, ensuring that we focused on sufficiently populated cell populations to enable reliable modeling of interactions. Next, we identified regions devoid of cells and incorporated this information as a spatial covariate within the model. By accounting for these regions, we hoped to avoid any bias in model fitting from assuming all areas in the spatial window were equally likely to contain cells. We further incorporated a spatial covariate of “distance to nearest blood vessel”, as a proxy of angiogenesis and oxygenation throughout the TME. An example of this spatial covariate can be seen in Figure 3.

**Figure 3:**
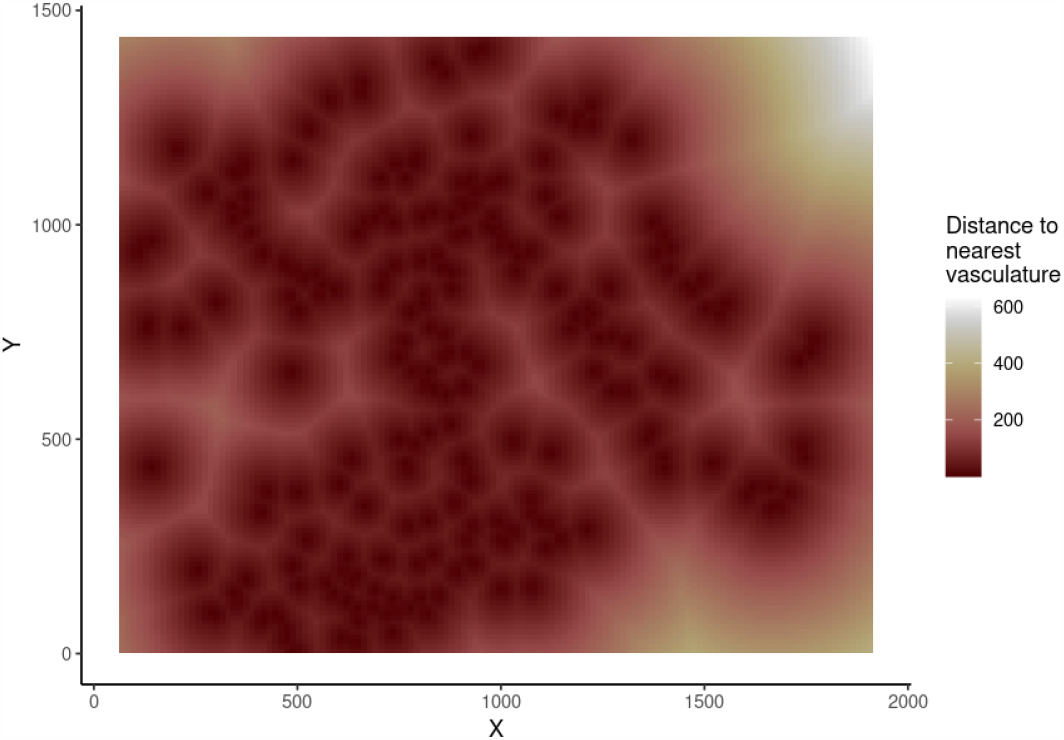
Spatial covariate of “distance to nearest blood vessel”, as imaged in spot 59_A.

The fitting of the MGPP models was performed using the ppjsdm package, an R package tailored for fitting saturated pairwise interaction Gibbs point process models [42]. Specifically, we employed an exponential interaction function to model the pairwise interactions between cell types at both short and medium ranges (that is, the functions *φ* and ψ in Equations 2.4 and 2.5). The interaction radii were set at 30, 70, and 150 microns, reflecting short-, medium- and long-range interaction distances, respectively. Lastly, 5,000 dummy points were included per cell type.

#### 2.3.2. Interpretation of Model Parameters and Predictions

After fitting the models, we extracted the fitted interaction parameters 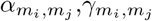 and *β* (corresponding to the effect that distance to nearest blood vessel had on the distribution of each cell type *m*_*i*_), along with the respective confidence interval for each parameter. We then plotted these parameters for one image as heatmaps, in order to demonstrate how to answer Questions 1 and 3. We also plotted the exponential potential functions scaled by each interaction parameter, in order to demonstrate how to answer Question 1a. We identified the cell type with the highest magnitude interaction parameter with tumor cells in each image, to answer Question 1b.

Next, we showed how one can answer Question 2 by identifying the cell type that has the largest magnitude contribution to the linear predictor of the conditional intensity of tumor cells at each point in space.

Lastly, we extracted the interaction parameters from all fitted models and plotted heatmaps of the difference in median interaction between the two patient groups, allowing us to answer Question 4.

#### 2.3.3. Heterogeneity in Model Parameter Estimates Within Patient Groups and Across Patients

In order to answer Question 5, we calculated the median absolute deviation (MAD) of each interaction parameter at two levels: pooled within each patient group, as well as across the whole patient cohort, and plotted the MAD estimates as heatmaps. This allows us to identify interactions that are highly variable and likely to be more contextdependent, as well as those that remain more consistent and homogeneous.

#### 2.3.4. Predictive Modeling of Patient Outcomes

We employed multivariate Cox proportional hazards models to estimate hazard ratios for interaction effects derived from the fitted MGPP models, in order to answer Question 6. Briefly, the interaction parameters for a given pair of cell types and spatial scale were extracted and used as a predictor of patient survival in a Cox model, while adjusting for the age and sex of each patient, as in Equation 2.6.

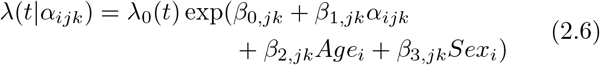

Here, *α*_*ijk*_ indicates the *α*_*jk*_ parameter from the *i*-th image. Models were fit for the *γ*_*jk*_ parameters in a similar fashion. Pairs of types that appeared in less than 10 images were excluded from this analysis. The resulting hazard ratios serve as potential prognostic biomarkers, indicating the prognostic significance of the identified spatial interactions within the tumor microenvironment. Furthermore, to account for the fact that there are multiple interaction estimates per patient (because there are multiple images, and thus multiple model fits, per patient), we fit these Cox models using a robust sandwich variance estimator for each fitted parameter that appropriately accounts for the correlation between interaction estimates within patients [45].

#### 2.3.5. Evaluation of Model Fit

Finally, in answering Question 7, we assessed the fit of the MGPP models by employing the Area Under the Curve (AUC) metric to quantify the predictive accuracy that the model had for each cell type within the tumor microenvironment. Briefly, the AUC is a metric that determines how well the predicted conditional intensity is able to partition the spatial window into regions of higher and lower concentrations of points [18]. By examining the AUC for each cell type, we are able to see how well the model predicts the spatial distribution of each cell type. We then calculated the median AUC per cell type, both within each patient group and across patient group, which helped identify cell types that could be reliably predicted by the MGPP models. Furthermore, we calculated the correlation between the AUC values and the abundance of each cell type, to clarify whether cell type abundance improved predictability of that cell type.

## 3. RESULTS FROM EXAMPLE ANALYSIS

### 3.1. Interpretation of Model Parameters and Predictions

We extracted the *β* parameters from each of the fitted models, and examples of the fitted parameters from one image are show in Table 1, allowing us to answer Question 3. The *β* parameters from this image demonstrate that the intensity of stroma, granulocytes and CD163+ macrophages are negatively associated with distance from vasculature that is, as the distance from vasculature increases, the less likely it is that stromal cells, granulocytes or CD163+ macrophages will be encountered.

**Table 1.**
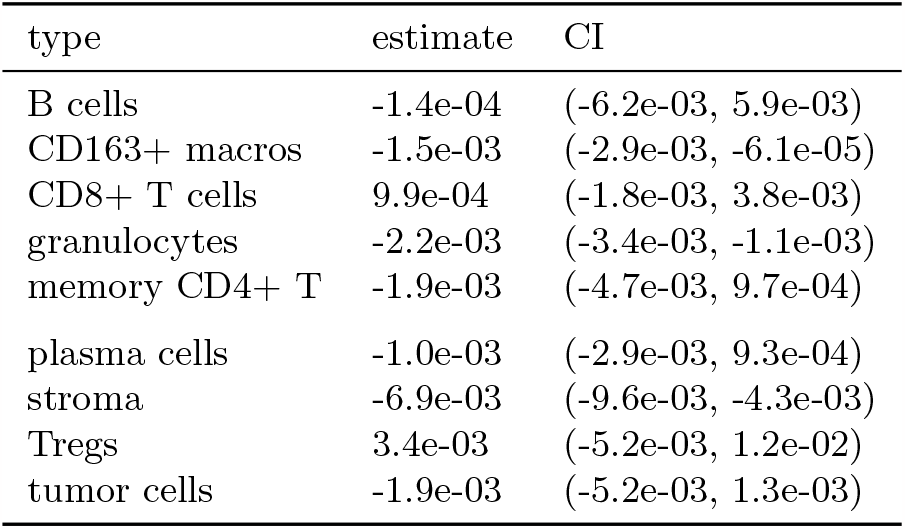
Coefficients 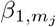 as estimated from spot 59_A.

We next extracted the 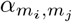and 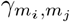 (subsequently *α* and *γ*) parameters from each of the fitted models. Examples of the fitted parameters from one image are given in Figure 4. The *α* parameters from this image demonstrate relatively consistent within-type positive interaction, or clustering, while between-type interactions at this scale tend to be negative, or repulsive. Of course, a significant outlier to this trend is the interaction between B cells and many other cell types, in particular with memory CD4+ T cells. The *γ* parameters, on the other hand, showcase a quite different story. Here, we again see that within-type interactions are generally positive (with the notable outlier of Tregs), though now tumor cells and CD163+ macrophages (or M2 macrophages) are generally positively associated with other cell types.

**Figure 4:**
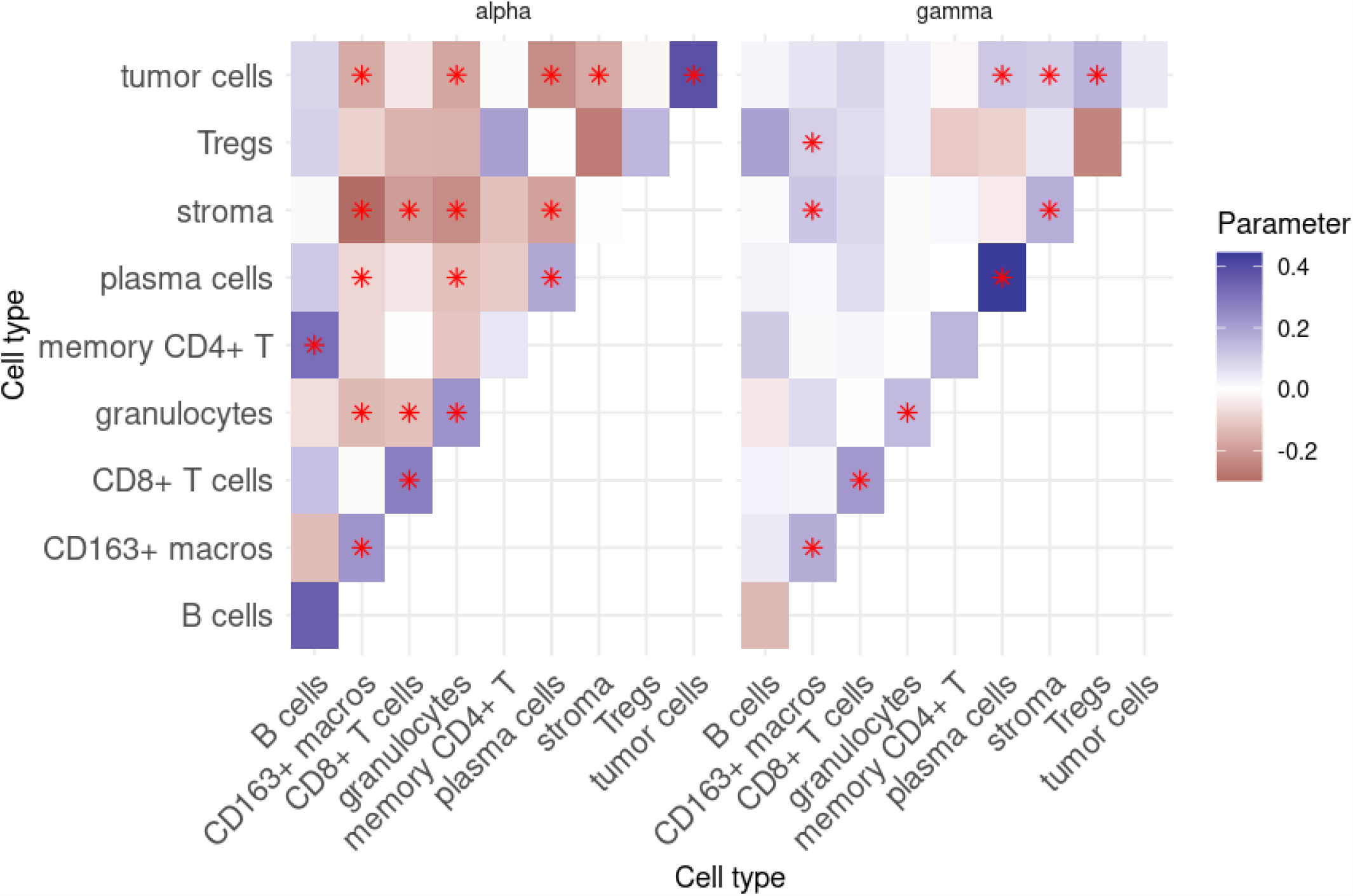
Interaction parameters 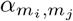and 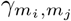 as estimated from spot 59_A. Stars indicate that the interaction parameter was found to be significant at a significance level of 0.05.

Heatmaps of the interaction parameters, as in Figure 4, allow one to quickly answer questions such as Question 1. From these heatmaps, we can also quickly answer questions like Question 1b: here, we can see that tumor cells seem to quite strongly positively correlated with each other at short scales, while quite negatively correlated with plasma cells, also at short scales. Interestingly, at larger scales, tumor cells are no longer correlated with each other, and are now positively correlated with plasma cells!

One is likely also to want to compare how these interaction parameters contrast between various patient groups, in order to answer questions like Question 4. Figure 5 demonstrate the difference in the median interaction parameters *α* and *γ* from each patient group (averaging across all images within a group). Strikingly, it appears that both the *α* and *γ* parameters for smooth muscle generally have the strongest differences in median interaction parameter, with patients from the CLR group generally exhibiting greater attraction between smooth muscle and Tregs, CD8+ T cells and CD163+ macrophages, while smooth muscle and granulocytes exhibit greater repulsion in CLR patients.

**Figure 5:**
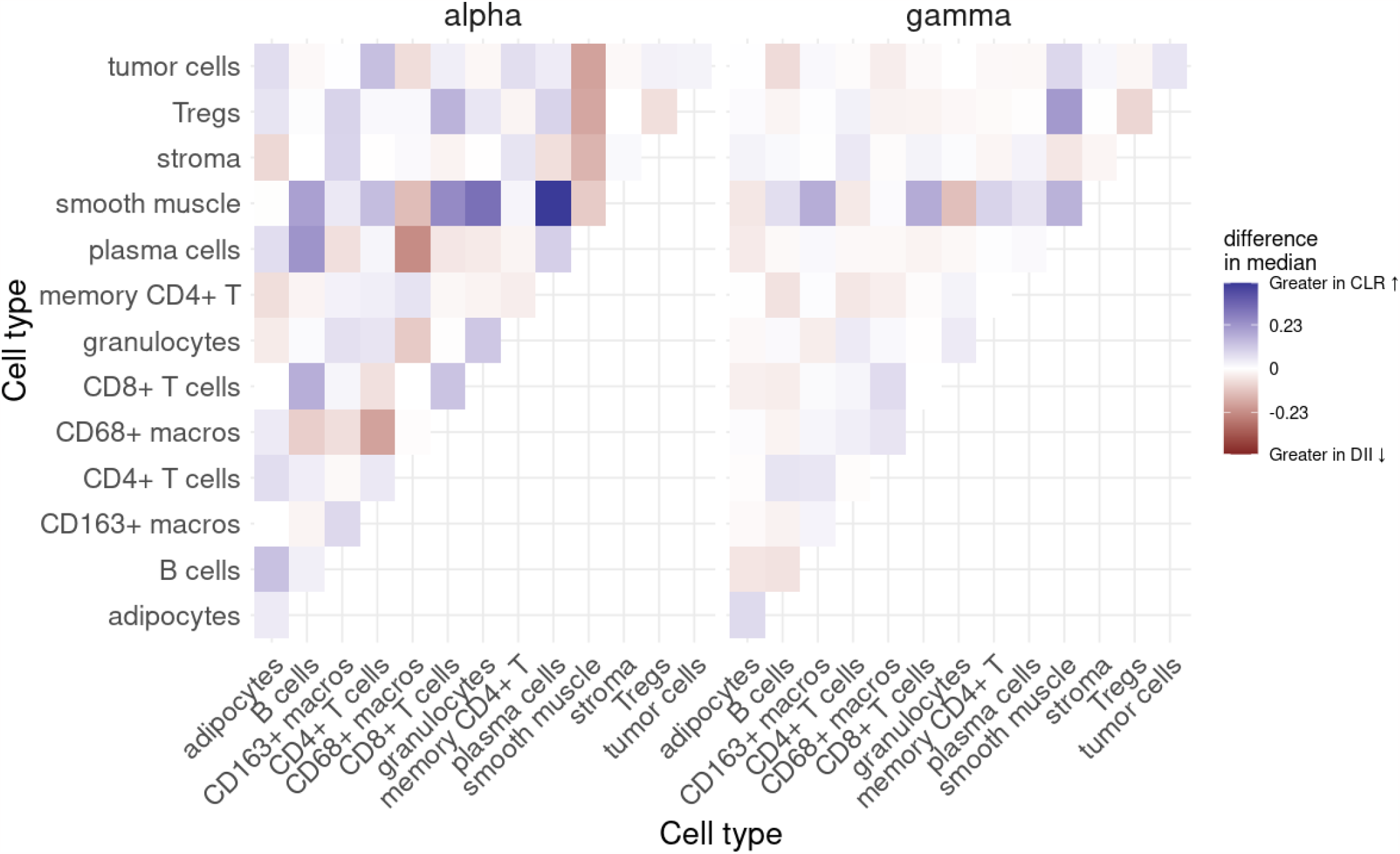
Difference in the median of the short-range interaction parameters 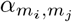 and 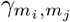 between patient groups.

Since the interaction parameters *α* and *γ* scale the potential functions used during the model fit (here, the exponential function), in order to answer questions liked Question 1a, it is also useful to view the pairwise potential functions themselves after they have been scaled by the appropriate parameters, as well as the combined effect of the short-range and medium-range scaled potential functions. An example of this can be seen in Figure 6, where the short-range potential (scaled by the appropriate *α*), medium-range potential (scaled by *γ*), and combined (sum of scaled short- and medium-range potentials) can be seen. The combined (or overall) potential thus gives a more complete picture between two cell types across spatial scales. Here we can see that B cells and memory CD4+ T cells are quite mutually attractive at shorter scales, but much less so at longer spatial scales. Such potentials can also be plotted simultaneously between all cell types of interest, as in Figure 7.

**Figure 6:**
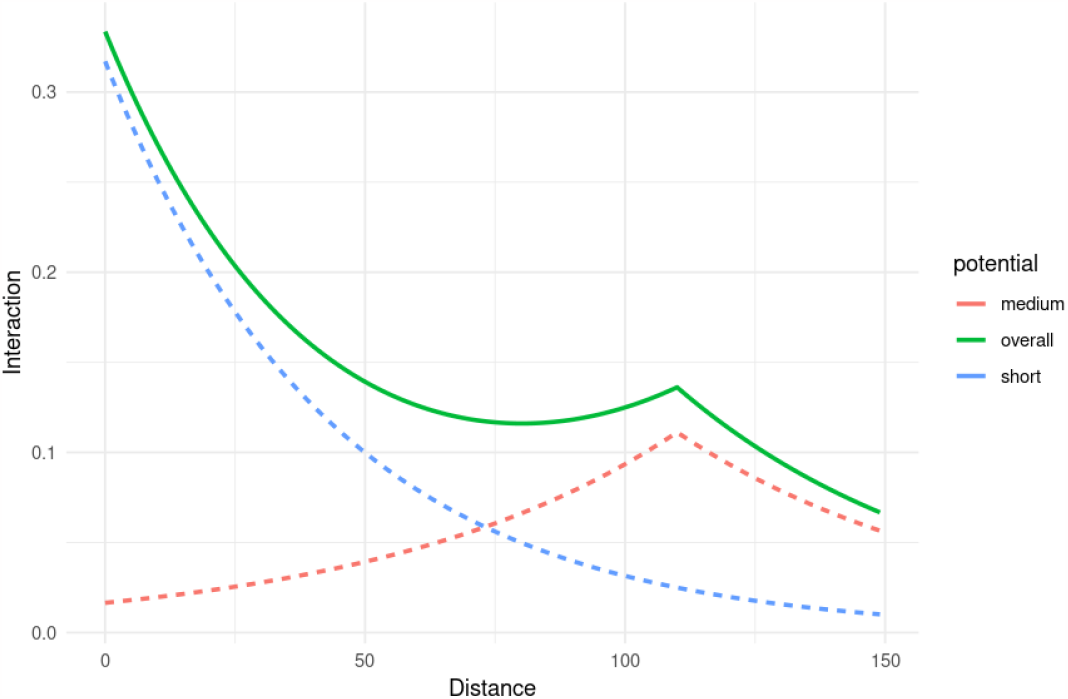
The short-range, medium-range and combined potentials between B cells and memory CD4+ T cells. From model fit on spot 59_A.

**Figure 7:**
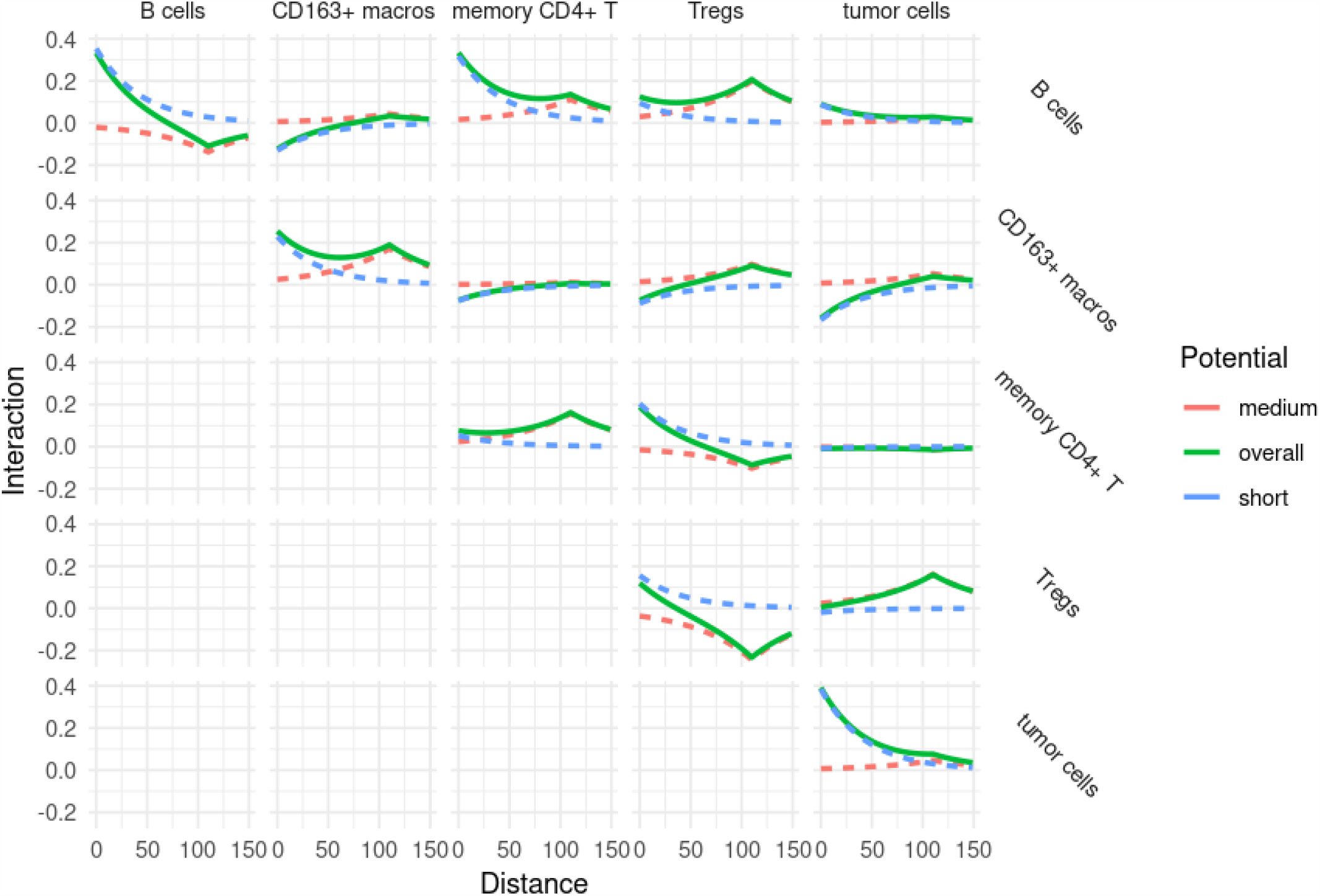
The short-range, medium-range and combined potentials between a subset of cell types. From model fit on spot 59_A.

From both of these figures, it is also apparent that there are many significant interactions occurring between these cell types, at multiple scales. These interactions indicate the spatial covariance between pairs of cell types that is unique to that pair. These quantities thus estimate the degree to which one cell type uniquely predicts the concentration of another, after accounting for the contributions to the prediction from all other cell types. From this information, we are thus able to assess which cell type most predicts the intensity of another cell type at each point in space, as in Figure 8. Such information allows one to answer questions like Question 2. Here we can see that different cell types are correlated more strongly with tumor cells at different points in the image. For example, granulocytes are the best predictors of tumor cells on the right side of the image, whereas CD163+ macrophages and stromal cells are the best predictors in other regions of the image.

**Figure 8:**
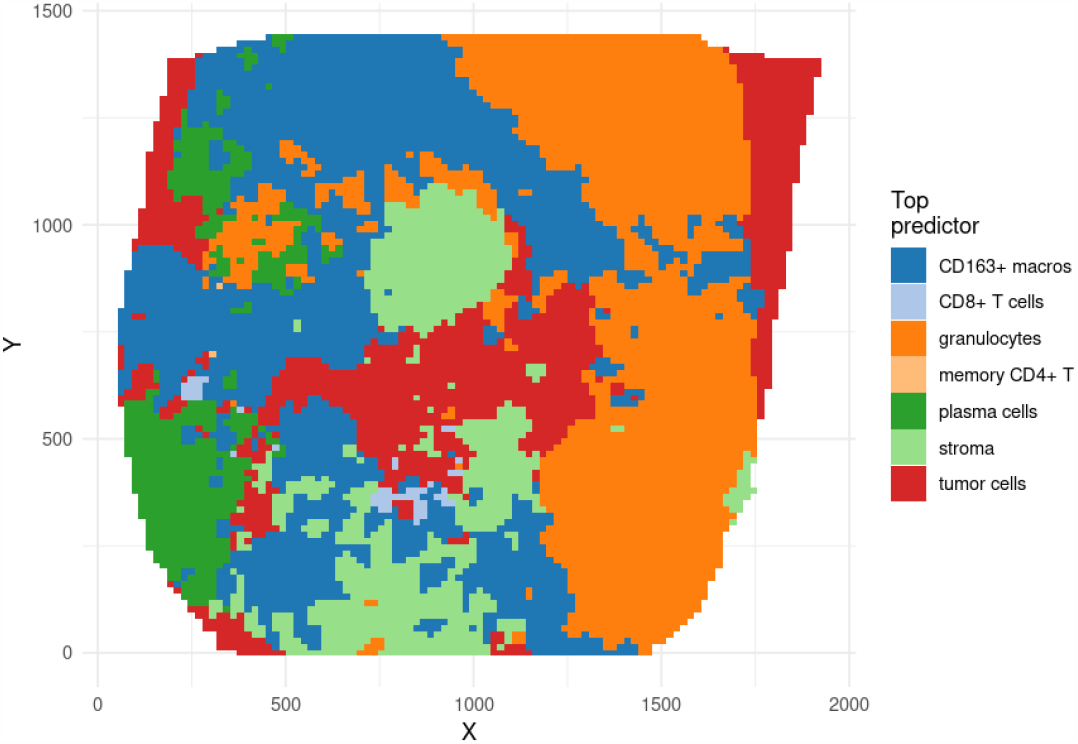
The top contributor to the predicted intensity of tumor cells. From model fit on spot 59_A.

### 3.2. Heterogeneity in Model Parameter Estimates Within Patient Groups and Across Patients

Estimates of the median absolute deviation of the shortrange interaction parameters *α* can be seen in Figures 9 and 10, allowing us to answer Question 5. In Figure 9, we can see that, across the entire patient cohort, within-type interactions for several of the immune cell types (in particular, B cells, CD163+ macros, CD4+ T cells, CD68+ macros and CD8+ T cells) are the most heterogeneous between images, while the interaction between smooth muscle and CD4+ T cells appears to be the most consistent across images.

**Figure 9:**
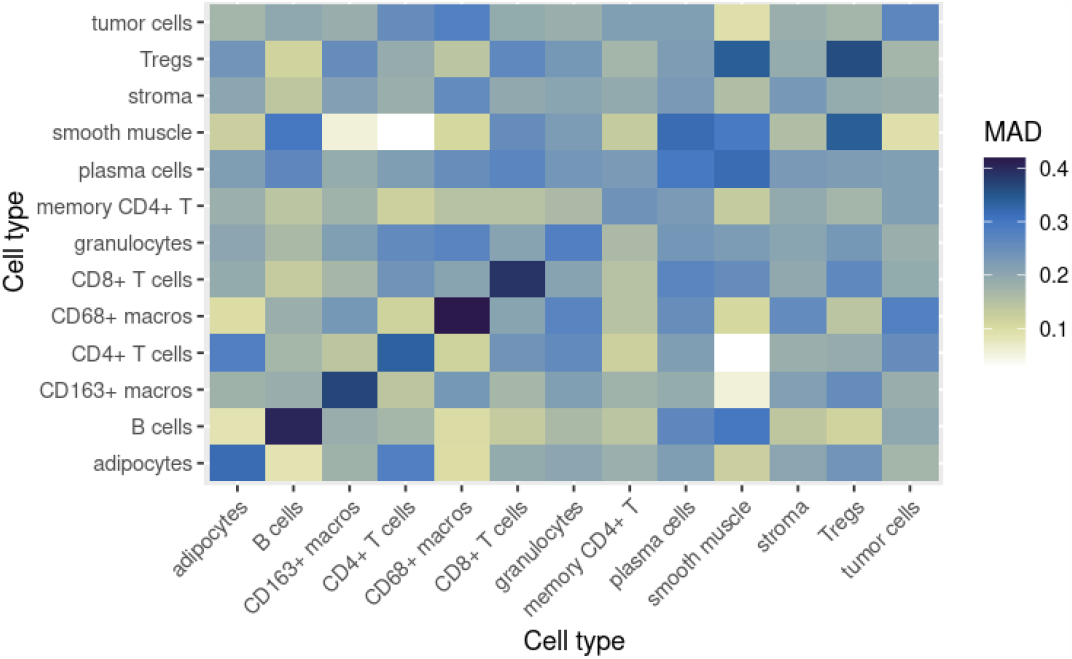
Estimates of the median absolute deviation (MAD) for *α* parameter estimates, across all patients.

**Figure 10:**
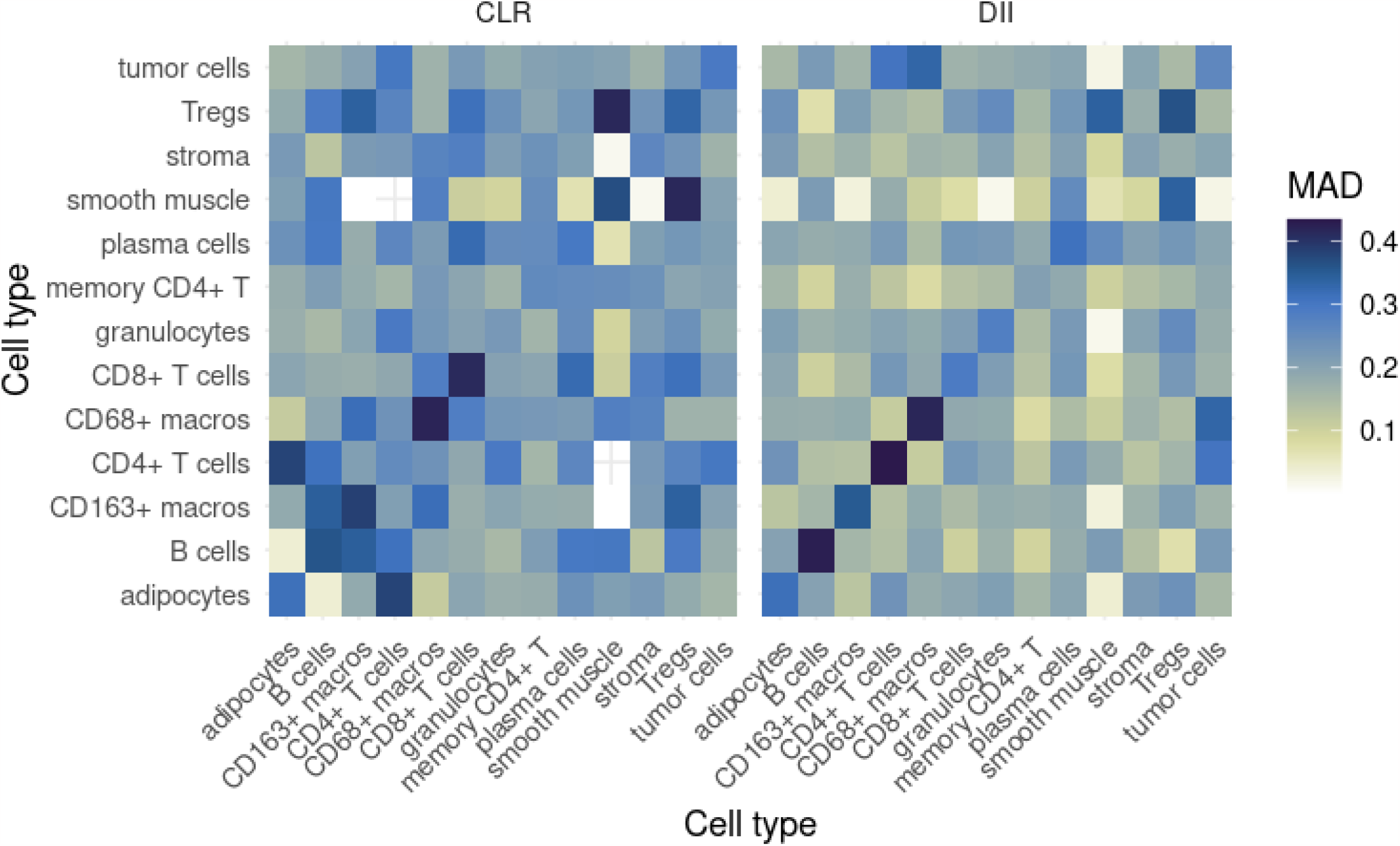
Estimates of the median absolute deviation (MAD) for *α* parameter estimates, in both patient groups.

In Figure 10, it appears that DII patients generally have more homogeneous interactions between images. However, interactions involving smooth muscle in CLR patients appear to be the most homogeneous among CLR patients, sometimes more so than their DII counterpart (for example, the interaction between smooth muscle and stroma). Interestingly, interactions involving smooth muscle also seem most different between the two patient groups, as detailed in the previous section.

### 3.3. Predictive Modeling of Patient Outcomes

Next, we turn our attention to answering Question 6. Estimated hazard ratios for both short- and medium-range interactions that were found to be significant after FDR adjustment for multiple hypothesis testing are shown in Figure 11. Conspicuously, increased short-range interactions between granulocytes and stroma seemed to be correlated with better patient outcomes, as well as granulocytes and M1 macrophages (CD68+ macros). Furthermore, the short-range interaction between M2 macrophages (CD163+ macrophages) and M1 macrophages seems to be indicative of poor patient outcome.

**Figure 11:**
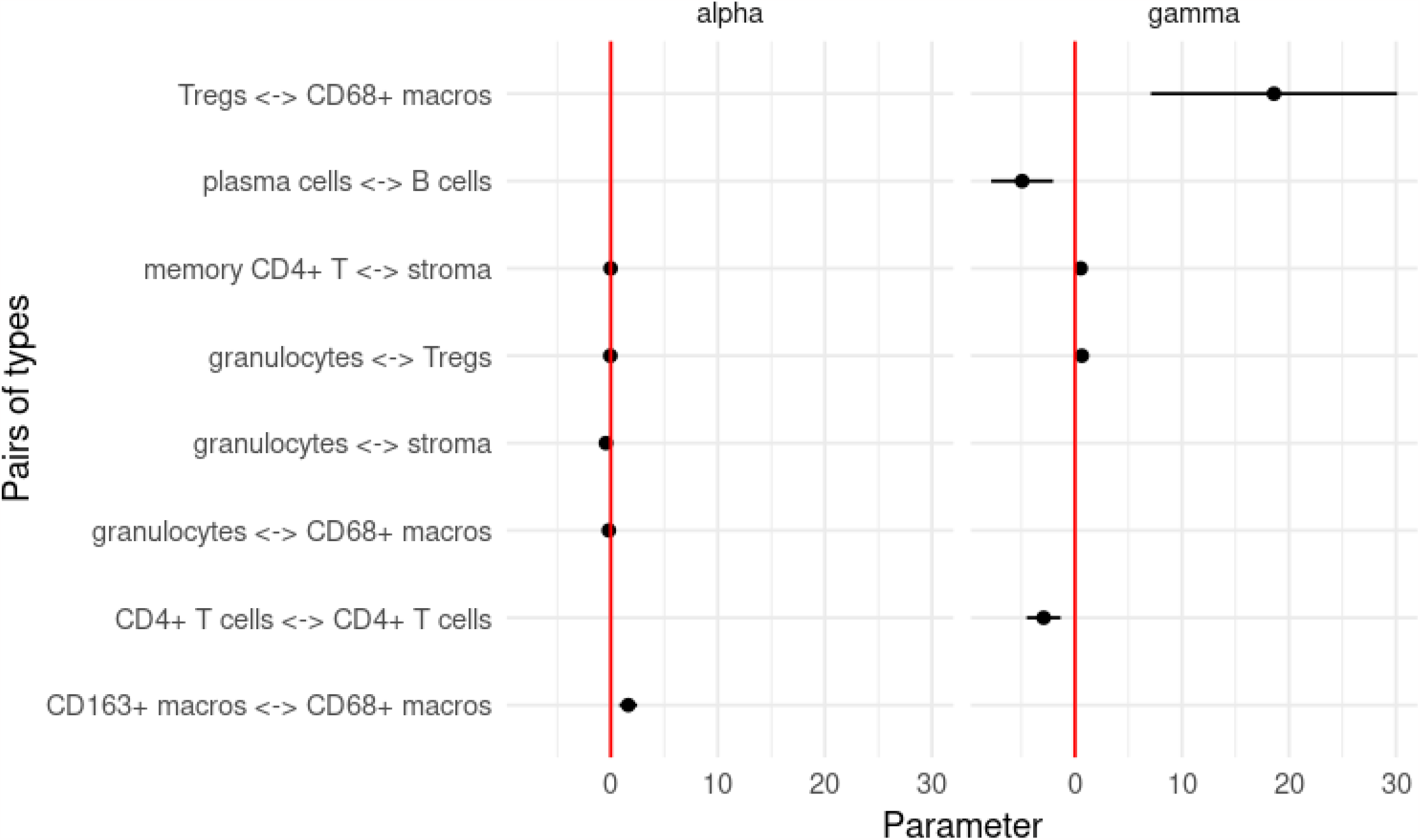
Log-hazard ratios estimated from short- and medium-range interaction parameters that were found to be significant after FDR adjustment.

**Figure 12:**
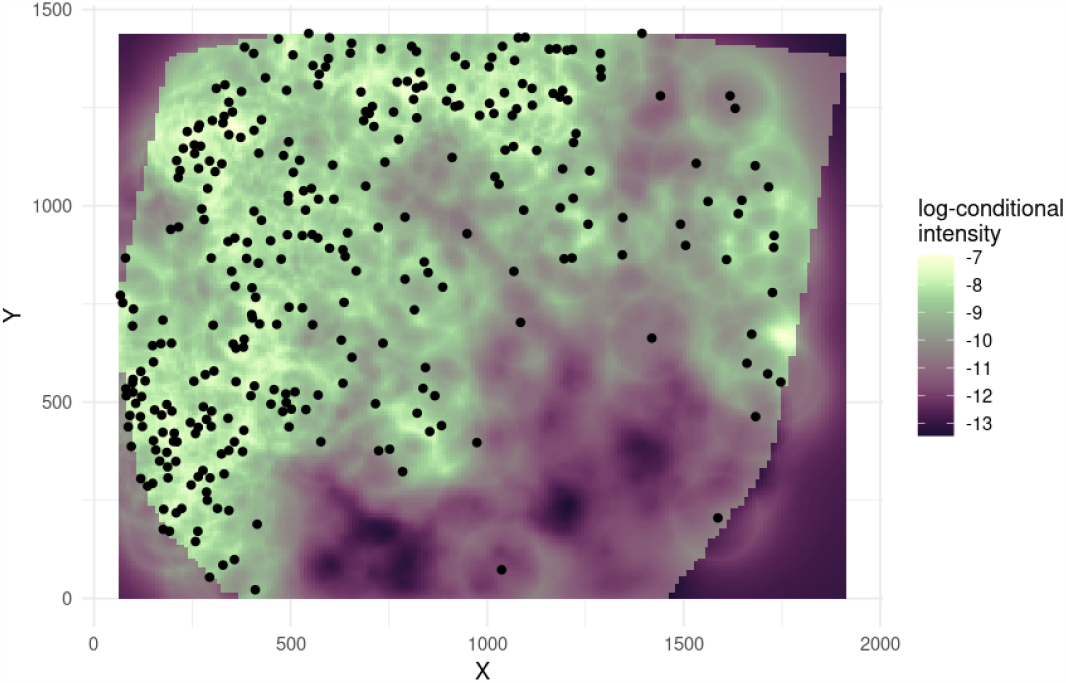
The predicted log-conditional intensity of plasma cells from the model fit for spot 59_A. The AUC for this cell type was 0.835, indicating high predictive ability for plasma cells.

At longer spatial scales (*γ* parameters from Figure 11), the Treg and M1 macrophage interaction seems to be highly correlated with poor patient outcome, while the plasma cell - B cell and CD4+ - CD4+ interactions were associated with better patient outcomes.

### 3.4. Evaluation of Model Fit

Lastly, we focus on the issue of model trust, in an effort to answer Question 7. An example of the predicted conditional intensity for plasma cells can be seen in Figure 12. The AUC (0.835) for this cell type seems relatively high, indicating that the model seems to predict the intensity of plasma cells quite well. Additionally, the observed positions of the plasma cells seem to match higher values of the predicted conditional intensity, mirroring the information given by the AUC.

Table 2 shows the median AUC per cell type across all images, as well as the median AUC within each patient group. From this table, we can see that the cell types tend to be similarly predictable on average. Furthermore, there does not seem to be a significant difference in predictability between the patient groups, except, notably, for CD4+ T cells. At first, it may seem that this could be simply because of the much larger number of CD4+ T cells in CLR patients than in DII patients. However, as noted in Table 3, B cells are represented almost 4 times more in CLR patients than DII patients, a larger order difference than found in CD4+ T cells, while maintaining median AUCs that are approximately the same.

**Table 2.**
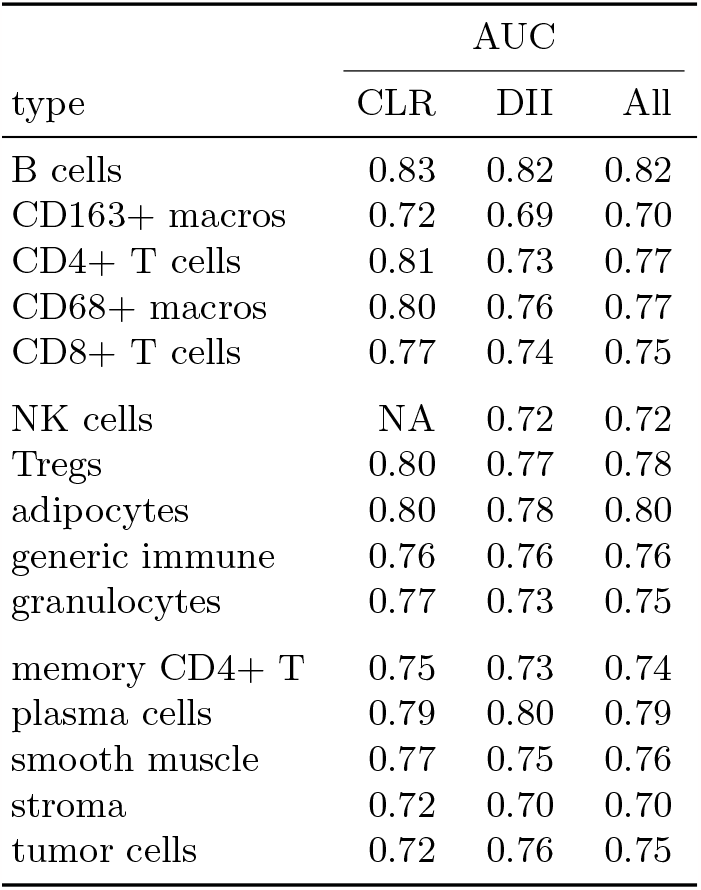
Median AUC for each cell type, across model fits from all images within each group and across the whole patient cohort.

**Table 3.** Abundance of each cell type in each patient group.

| type | abundance |  |
| --- | --- | --- |
|  | CLR | DII |
| adipocytes | 8.5 | 4.0 |
| B cells | 14.0 | 17.0 |
| CD163+ macros | 172.5 | 316.5 |
| CD4+ T cells | 5.5 | 2.0 |
| CD68+ macros | 9.5 | 20.0 |
| CD8+ T cells | 62.0 | 109.0 |
| generic immune | 12.0 | 9.0 |
| granulocytes | 27.0 | 123.5 |
| memory CD4+ T | 53.0 | 80.0 |
| NK cells | 1.5 | 1.0 |
| plasma cells | 28.0 | 22.0 |
| smooth muscle | 77.0 | 124.0 |
| stroma | 104.0 | 157.5 |
| Tregs | 8.0 | 21.0 |
| tumor cells | 102.5 | 222.0 |
| vasculature | 61.5 | 75.0 |

To make the aforementioned precise, we measured the correlation between the AUC and abundance of each cell type. Table 4 shows the correlation between cell type and abundance, as quantified using AUCS from all model fits. All correlations can be seen to be negative, some with a quite large magnitude. This indicates that simply having more cells of a given type in an image does not make that cell type necessarily easier to predict using MGPP models. This likely indicates the importance of incorporating higher-order information on cell-type interactions, beyond cell-type abundance, in order to correctly predict distributions of cells.

**Table 4.** Correlation between AUC score and abundance for each cell type.

| type | correlation |
| --- | --- |
| B cells | -0.18 |
| CD163+ macros | -0.39 |
| CD4+ T cells | -0.30 |
| CD68+ macros | -0.41 |
| CD8+ T cells | -0.49 |
| NK cells | -0.99 |
| Tregs | -0.35 |
| adipocytes | -0.42 |
| generic immune | -0.59 |
| granulocytes | -0.54 |
| memory CD4+ T | -0.29 |
| plasma cells | -0.41 |
| smooth muscle | -0.58 |
| stroma | -0.49 |
| tumor cells | -0.16 |

Overall, from Table 2, we can see that the models predict the spatial distribution of cells quite well, indicating that we can likely trust the results from our models.

### 3.5. Comparison with Previous Analysis

Since the CRC dataset used here comes from a previously published study [16], here we outline a few key points in comparison to the results from that study.

1. There is evidence of greater attraction between tumor cells and some subsets of immune cells in DII patients - for example, B cells and M1 macrophages - than CLR patients - similarly, [16] identified the presence of a mixed tumor/immune compartment in DII patients, as opposed to CLR patients, in whom the tumor and immune components were more segregated. Interestingly, though, at short ranges, CD4+ T cells and tumor cells are found to be more attractive in CLR patients (Figure 5).
2. Schurch et al found much greater contact between a T-cell enriched neighborhood and a macrophage-enriched neighborhood in DII patients. Similarly, we found that M1 macrophages were more attractive with CD4+ T cells and plasma cells in DII patients at short ranges (Figure 5).
3. The authors further identified that the functional state of a granulocyte-enriched cellular neighborhood was indicative of patient survival in DII patients. Similarly, we also found that granulocyte associations were frequently implicated as the most prognostic biomarkers across patient groups derived from our models (Figure 11).

## 4. DISCUSSION

In this paper, we apply a novel approach for analyzing ecological interactions within the tumor microenvironment (TME). Specifically, we employ a multitype Gibbs point process model (MGPP) as a joint species distribution model, tailored to the complexities of the TME. This modeling framework was previously introduced as a statistical analysis framework in forest ecology [42]. Our aim in this work is to highlight the significance of this modeling framework and its potential implications for addressing key questions in cancer biology.

Our primary contribution lies in the application of a specific MGPP model to characterize the data-generating process governing cell-cell and cell-environment interactions in the TME. Given the intricate nature of the TME, where diverse cell types interact spatially, the MGPP framework provides a powerful tool for capturing and quantifying these interactions. What sets this modeling framework apart from many others that currently exist is its ability to model relationships between cell types simultaneously, while also being able to quantify the uncertainty in these associations as well as produce predictions. Researchers currently lack such a unified statistical framework capable of jointly modeling interactions between cell types and spatial variables across multiple spatial scales. With a specific MGPP model, we showcase how to bridge this gap, enabling a diverse array of inquiries.

Our emphasis extends beyond model development to model validation and diagnostics. We highlight the utility of our fitted models in generating predictions in order to examine goodness-of-fit and demonstrate the area under the curve (AUC) as a concise summary of model fit. Additionally, we stress the availability of further model diagnostics based on residuals.

Beyond model fitting, our study demonstrates the utility of extracting estimated parameters from single-image models and summarizing them across patient cohorts. This approach can serve as a valuable tool for identifying potential prognostic biomarkers, furthering our understanding of cancer biology and personalized medicine.

While hyperparameter selection is a critical aspect of model development, we do not delve deeply into this topic in this paper. Interested readers can find comprehensive discussions on the selection of hyperparameters, including the scale and shape of potential functions, the saturation parameter and the number of dummy points, in [42].

Lastly, MGPP models offer a unique capability which we also did not address in this paper: the generation of simulations while conditioning on various aspects of the data. This feature has the potential to serve two primary purposes—validating model outputs by comparing them to expected results and simulating new cell types based on different configurations. Such simulations can shed light on the spatial organization of cell types within the TME.

In conclusion, our investigation into cell-type interactions in the tumor microenvironment using multitype Gibbs point process models presents a valuable tool for researchers interested in the complex ecology of the TME. This framework empowers researchers to explore complex questions, validate models, and gain deeper insights into the dynamics of the TME, with potential implications for cancer research and therapy.

## CONFLICTS OF INTEREST

A.R serves as member for Voxel Analytics, LLC and consults for Telperian Inc. and Tempus Labs Inc.

## FUNDING

This work was supported by the Advanced Proteogenomics of Cancer training grant (T32 CA140044), an R37 MERIT award from the NIH (R37 CA214955-01A1), an NSF grant (DMS-2152776) and MICDE CAtalyst grants.

